# Cell-state dependent regulation of PPAR_γ_ signaling by ZBTB9 in adipocytes

**DOI:** 10.1101/2024.03.04.583402

**Authors:** Xuan Xu, Alyssa Charrier, Sunny Congrove, David A. Buchner

## Abstract

Adipocytes play a critical role in metabolic homeostasis. Peroxisome proliferator-activated receptor-_γ_ (PPAR_γ_) is a nuclear hormone receptor that is a master regulator of adipocyte differentiation and function. ZBTB9 was predicted to interact with PPAR_γ_ based on large-scale protein interaction experiments. In addition, GWAS studies in the type 2 diabetes (T2D) Knowledge Portal revealed associations between Z*btb9* and both BMI and T2D risk. Here we show that ZBTB9 positively regulates PPAR_γ_ activity in mature adipocytes. Surprisingly Z*btb9* knockdown (KD) also increased adipogenesis in 3T3-L1 cells and human preadipocytes. *E2F* activity was increased and E2F downstream target genes were upregulated in *Zbtb9*-KD preadipocytes. Accordingly, RB phosphorylation, which regulates E2F activity, was enhanced in *Zbtb9*-KD preadipocytes. Critically, an E2F1 inhibitor blocked the effects of *Zbtb9* deficiency on adipogenic gene expression and lipid accumulation. Collectively, these results demonstrate that *Zbtb9* inhibits adipogenesis as a negative regulator of *Pparg* expression via altered RB-E2F1 signaling. Our findings reveal complex cell-state dependent roles of ZBTB9 in adipocytes, identifying a new molecule that regulates adipogenesis and adipocyte biology as both a positive and negative regulator of PPAR_γ_ signaling depending on the cellular context, and thus may be important in the pathogenesis and treatment of obesity and T2D.

## Introduction

Obesity is a one of the greatest health challenges facing our world today. Comorbidities include type 2 diabetes (T2D), cardiovascular disease, and certain types of cancer which collectively increase the risk of premature mortality (1). Treatments for these metabolic diseases include surgery, lifestyle modifications, and therapeutics such as the recently developed incretin mimetics (2). However, despite tremendous recent successes in the treatment of obesity and T2D, there remain many individuals for which safely maintaining a healthy body weight and long-term glucose homeostasis remains challenging such that new therapeutic approaches are a major unmet clinical need (3,4). Adipose tissue is the body’s primary site of fat storage and has a vital role for integrating and communicating metabolic signals. Adipose tissue expands when energy intake exceeds energy expenditure, storing the excess nutrients in lipid droplets as inert triacylglycerol (TAG) and thereby maintaining metabolic homeostasis (5). Adipocytes within adipose tissues store excess energy via hyperplasia, an increase in the number of adipocytes, and hypertrophy, an increase in the size of adipocytes (6). Discovering the mechanisms regulating adipocyte function and adipocyte differentiation is central for understanding the pathophysiology of obesity and its related metabolic diseases and identifying new therapeutic opportunities (7–9).

Peroxisome proliferator-activated receptor _γ_ (PPAR_γ_) is a member of the nuclear hormone receptor family of ligand-inducible transcription factors (10). PPAR_γ_ is a master transcriptional regulator of adipogenesis and also a key regulator of gene expression in mature adipocytes (11,12). Thiazolidinedione (TZDs), synthetic ligands of PPAR_γ_, are activators of PPAR_γ_ with robust insulin-sensitizing activities (13). Although treatment with these compounds causes weight gain and an array of additional safety issues (e.g., cardiovascular risk and fluid retention), TZDs were once widely used to treat T2D (14). A more complete understanding of PPAR_γ_ activation and signaling will potentially lead to new and improved therapies for T2D. Control of PPAR_γ_ transcriptional activity depends on multi-protein complexes containing dimerization partners and other co-regulators, which each have their own physiological effects on insulin resistance and metabolic dysfunction (15–17). PPAR-gamma coactivator-1_α_ (PGC-1_α_) is a PPAR_γ_ coactivator which serves as a scaffolding protein to recruit a variety of other coactivators. PGC-1_α_ regulates the activity of PPAR_γ_ on adaptive thermogenesis and fatty acid oxidation by interacting with the PPAR_γ_/RXR_α_ heterodimer. This interaction stimulates the expression of UCP-1, resulting in enhanced metabolic rate and insulin sensitivity, and resistance to obesity (18–21). NCoR and SMRT are other examples of well characterized PPAR_γ_ co-regulators that function to recruit histone deacetylase (HDAC) complexes, which covalently modify nucleosomes to compact DNA and repress transcription (22). In the absence of ligand, PPAR_γ_ recruits the transcription corepressors NCoR and SMRT to downregulate PPAR_γ_-mediated transcriptional activity. Gene silencing of *Ncor* or *Smrt* in 3T3-L1 preadipocytes increases adipocyte differentiation (22–24). These cofactors, and many others, have specific physiological functions and differential effects on regulating the transcriptional action of PPAR_γ_, and therefore unique non-redundant roles in lipid and energy metabolism.

Zinc finger protein 407 (ZFP407) was first identified as a positive regulator of insulin-stimulated glucose uptake via regulation of the PPAR_γ_ signaling pathway both *in vitro* and *in vivo* (25,26). ZFP407 was subsequently shown to directly interact with the PPAR_γ_/RXRα protein complex (27). To better understand the mechanism by which PPAR_γ_ activity is regulated, we focused on discovering novel proteins that interact with the PPAR_γ_ and ZFP407 transcriptional regulatory protein complex. Towards this end, a previously performed high-throughput mammalian two-hybrid analysis of transcription factor protein-protein interactions (28) identified, among many other novel interactions, that ZBTB9 directly interacted with multiple proteins critical for PPAR_γ_ signaling, including PPAR_γ_ itself, but also including ZFP407 and PGC-1β (26,29–32). Furthermore, *Zbtb9* is the closest gene to a series of SNPs that are significantly associated with Body Mass Index (BMI). The lead SNP among them, *rs11757081*, was significantly associated with BMI (p < 10^-15^) in a GWAS meta-analysis of over 3 million individuals (33). *Zbtb9* is also the closest gene to SNP *rs210192*, which was significantly associated with T2D risk (p = 4.52e^-8^) in a similar meta-analysis (33). These data suggest that ZBTB9 may play a critical role in determining an individual’s risk of developing metabolic disease and T2D.

To assess the potential role of ZBTB9 in PPAR_γ_ signaling and adipocyte biology, we generated *Zbtb9*-deficienct mouse and human adipocytes and preadipocytes, and demonstrated for the first time the critical role ZBTB9 has on adipogenesis and insulin sensitivity via modulation of PPAR_γ_ signaling. We further show that ZBTB9 is a negative regulator of early steps in adipogenesis via E2F-dependent regulation of *Pparg* expression. Interestingly, in mature adipocytes, ZBTB9 is required for PPAR_γ_ signaling and insulin response via a different mechanism that regulates PPAR_γ_ activity, but not expression. Together, our results provide the first functional characterization of ZBTB9 in adipocytes, revealing unique cell-state-dependent regulation of PPAR_γ_ signaling in both early adipogenesis and mature adipocyte function.

## Results

### ZBTB9 positively regulates PPAR*_γ_* activity in mature adipocytes

A high throughput mammalian two-hybrid analysis of transcription factor protein-protein interactions suggested that ZBTB9 interacts with PPAR_γ_ and other PPAR_γ_-interacting proteins including ZFP407 (28). To further test whether ZBTB9 interacts with these proteins, a co-IP was performed in differentiated 3T3-L1 adipocytes. The co-IP confirmed that ZBTB9 does indeed interact with the PPAR_γ_/RXRα/ZFP407 complex (Fig. 1A). To investigate the role of ZBTB9 in PPAR_γ_ signaling, ZBTB9 and PPAR_γ_ were co-expressed with a PPAR_γ_ response element (PPRE) luciferase reporter plasmid that contains 3 copies of the canonical PPAR_γ_ DNA binding sequence DR1. Together, ZBTB9 and PPAR_γ_ increased expression from the PPAR_γ_ reporter construct (Fig. 1B-C), suggesting that ZBTB9 positively regulates PPAR_γ_ activity through the canonical PPAR_γ_ response element.

**Figure 1.**
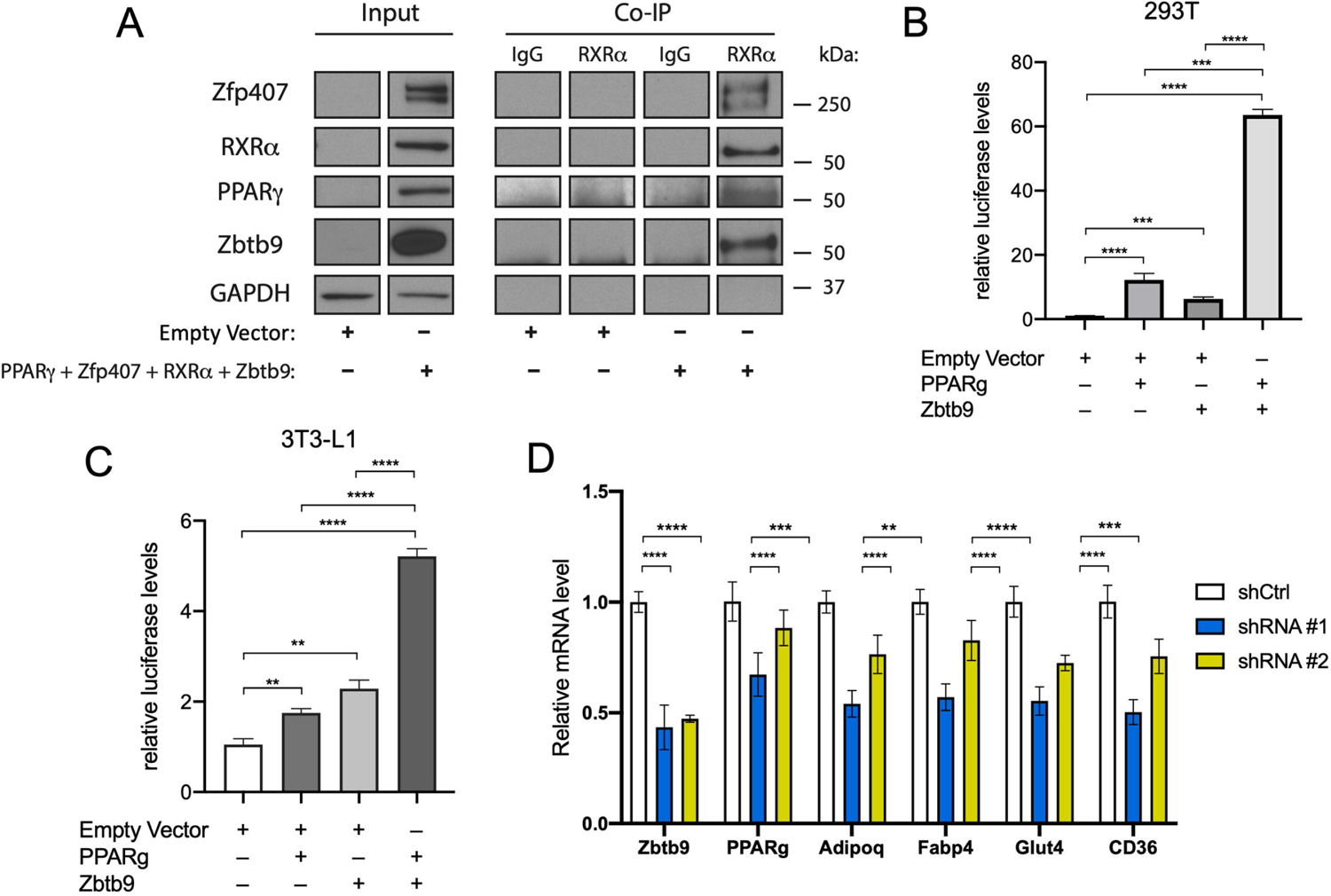
ZBTB9 is a positive regulator of PPAR_γ_ signaling in mature adipocytes. (A) Co-IP using anti-RXRα antibody of exogenously expressed proteins as indicated in differentiated 3T3-L1 cells. (B) PPRE luciferase reporter activity from 293T cells or from (C) differentiated 3T3-L1 adipocytes transfected with an empty vector, *Zbtb9* cDNA, *Pparg* cDNA as indicated. (D) *Zbtb9* was knocked down in differentiated 3T3-L1 adipocytes with 2 independent shRNAs (shRNA #1, shRNA #2) and compared to the control shRNA (shCtrl). Gene expression was measured by qRT-PCR. ** p < 0.01, *** p < 0.001, *** p < 0.0001. n=3-5/group.

To explore the role of ZBTB9 on PPAR_γ_ target gene expression under more physiological conditions, we reduced *Zbtb9* expression by lentivirus-mediated shRNA in differentiated 3T3-L1 adipocytes, and observed consistently decreased expression of a number of well-characterized PPAR_γ_ target genes in *Zbtb9* knockdown (KD) cells compared to the control (Fig. 1D). Altogether, this data suggests that ZBTB9 acts as a positive regulator of PPAR_γ_ signaling in adipocytes.

### Zbtb9 is a negative regulator of adipogenesis

Given the interaction between PPAR_γ_ and ZBTB9 (Fig. 1A), the role of ZBTB9 in regulating PPAR_γ_-dependent gene expression in mature adipocytes (Fig. 1B-D), and the central role of PPAR_γ_ in adipogenesis (31), we next examined whether ZBTB9 played a role in adipocyte differentiation. Two independent shRNAs were used to inhibit *Zbtb9* expression in 3T3-L1 preadipocytes. The cells were then induced to undergo adipocyte differentiation in the context of reduced *Zbtb9* expression. Surprisingly, lipid accumulation, as assessed by Oil Red O staining, was significantly increased in the *Zbtb9* KD cells relative to control cells (Fig. 2A-B). Consistent with the lipid accumulation, expression of the adipogenic genes *Pparg*, *Adipoq*, *Fabp4* and *Glut4* were all significantly higher in *Zbtb9*-KD cells relative to control cells (Fig. 2C). The increased levels of *Pparg* and *Fabp4* expression in *Zbtb9*-KD cells were observed at different time points during differentiation, starting within three days of inducing adipogenesis, with bigger effects at day 3 and day 7, suggested that ZBTB9 plays an early role in regulating key genes that are critical to the molecular induction of adipogenesis (Fig. 2D).

**Figure 2.**
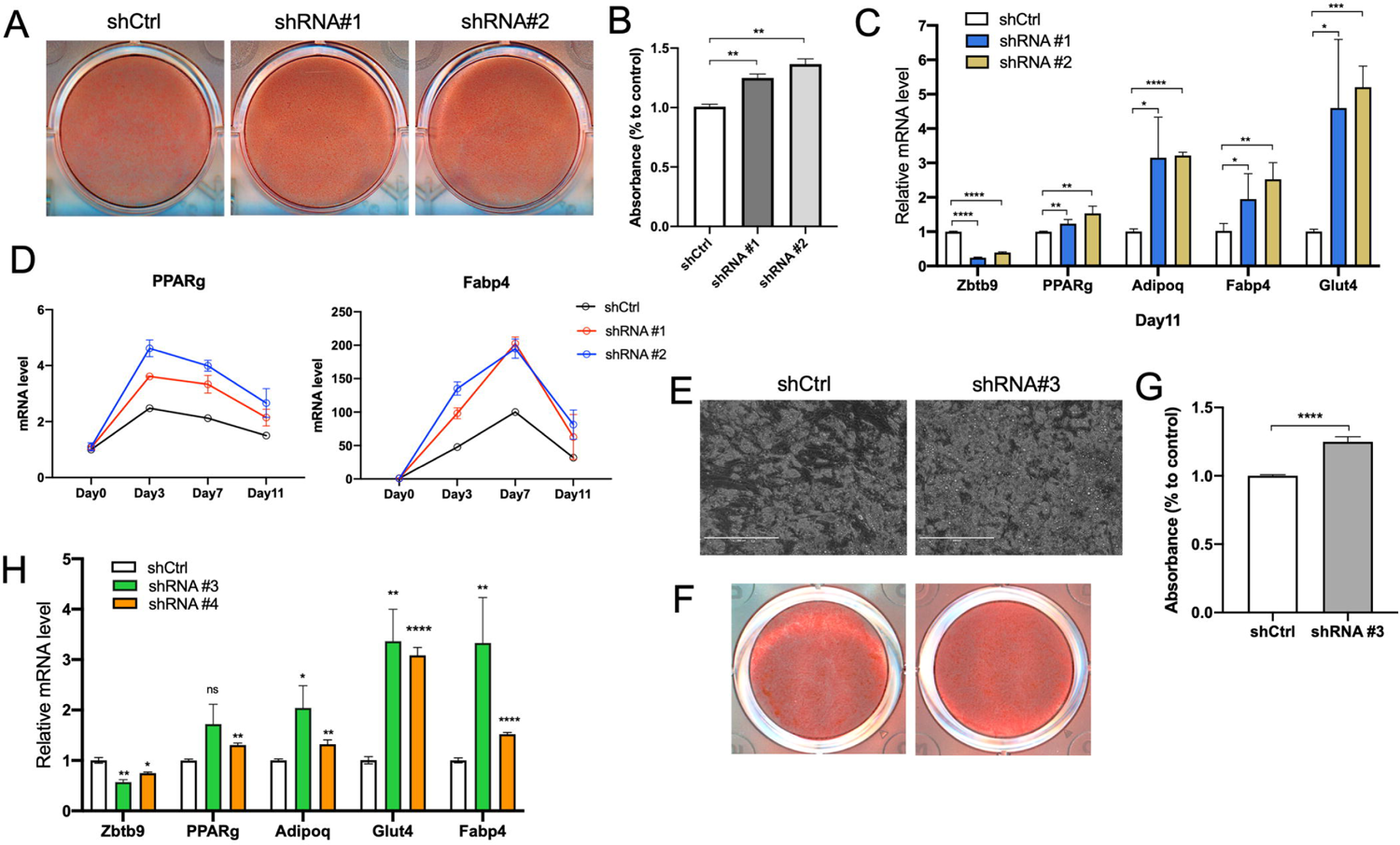
*Zbtb9* deficiency increases adipogenesis. (A) Oil Red O staining and (B) quantification to assess lipid accumulation in *Zbtb9*-KD 3T3-L1 cells or control cells. (C) *Zbtb9* and adipogenic gene expression at the end of differentiation (day 11) in *Zbtb9*-KD 3T3-L1 cells or control cells as determined by qRT-PCR. (D) Time course of *Pparg* and *Fabp4* gene expression during differentiation at D0, D3, D7 and D11 in *Zbtb9*-KD 3T3-L1 cells or control cells as determined by qRT-PCR. (E) Lipid accumulation in differentiated human adipocytes treated with *hZBTB9* shRNA (shRNA #3) or control shRNA (shCtrl) shown with brightfield images and (F) oil red O staining and (G) oil red O quantification. (H) *ZBTB9* was knocked down with 2 independent shRNAs (shRNA #3, shRNA #4) in human preadipocytes. Gene expression was measured by qRT-PCR at the end of differentiation (day 14). * p < 0.05, ** p < 0.01, **** p < 0.0001, ns: not significant. n=3-5/group.

ZBTB9 protein is highly conserved between mice and humans (77% identical at the amino acid level, Fig. S1). The level of conservation is even higher specifically within the BTB domain and the two C2H2 zinc finger domains (96% identical, Fig. S1). Thus, to extend the findings observed in mouse 3T3-L1 cells, and test whether ZBTB9 has a similar function during adipogenesis in humans, analogous studies were performed in a human-derived cell model of adipogenesis. *ZBTB9* levels were reduced by lentiviral-mediated shRNA treatment in human preadipocytes, and upon adipogenesis, there was again a significant increase in both lipid accumulation, as assessed by oil red O staining (Fig. 2E-G) and adipogenic marker gene expression (Fig. 2H). Together, these data suggest that while ZBTB9 positively regulates PPAR_γ_ activity in mature adipocytes, it regulates the early stages of adipogenesis by an alternative mechanism in a manner that is evolutionarily conserved in both humans and mice.

### Transcriptome-wide profiling reveals that Zbtb9 broadly regulates the expression of genes within the adipogenesis pathway

To identify the genes and biological processes regulated by *Zbtb9* during differentiation, *Zbtb9*-KD 3T3-L1 cells were analyzed by RNA-Seq throughout adipocyte differentiation, beginning at day 0 (before adipogenic induction), as well as days 3, day 7, and finally day 11 when the cells are terminally differentiated into adipocytes. Principle component analysis (PCA) was performed to test whether samples clustered with each other at each time point during adipogenesis. Figure 3A shows the results of the PCA, demonstrating that samples at day 0 and day 3 were distinct from those at day 7 and day 11. On day 0 or day 3, there was a clear distinction between KD and control samples, whereas samples at day 7 and day 11 tended to cluster together, regardless of whether the cells were treated with a control or *Zbtb9*-targeting shRNA. This suggests that the cells have different gene profiles at later time points compared to early ones during adipocyte differentiation, consistent with the transition from preadipocytes to adipocytes. Additionally, bigger differences were observed between KD and control samples at day 7 and day 11, respectively, as compared with early time points, suggesting bigger effects of *Zbtb9* on the gene profiles later during differentiation. Differentially expressed genes (DEGs) were identified at each time point during differentiation (FDR ≤ 0.05). In general, more DEGs were observed upon adipogenic differentiation, consistent with the PCA results. A total of 826, 1103, 4088 and 3195 genes were found to be differentially regulated, with 352, 594, 2198 and 1639 gene transcripts upregulated, and 474, 509, 1890 and 1556 genes downregulated at day 0, day 3, day 7 or day 11 respectively (Fig. 3B-D). Gene set enrichment analysis showed that upregulated genes were enriched for lipid metabolism-related pathways, and included adipogenesis, oxidative phosphorylation, fatty acid metabolism, and MTORC1 signaling (Fig. 3E). Increased expression of most of the pathways in *Zbtb9*-KD cells relative to control cells was observed beginning at day 3. These expression differences are likely to be largely attributable to the upregulation of *Pparg* in KD cells compared with control cells that was also seen at this time point during differentiation (Fig. 2D and Fig. 4). Significantly upregulated genes in KD vs. control cells of adipogenesis pathway at days 3, 7, and 11 included many well-known adipogenic and *Pparg* target genes including *Adipoq*, *Fabp4*, *Cd36*, and *Lpl* (Fig. 4).

**Figure 3.**
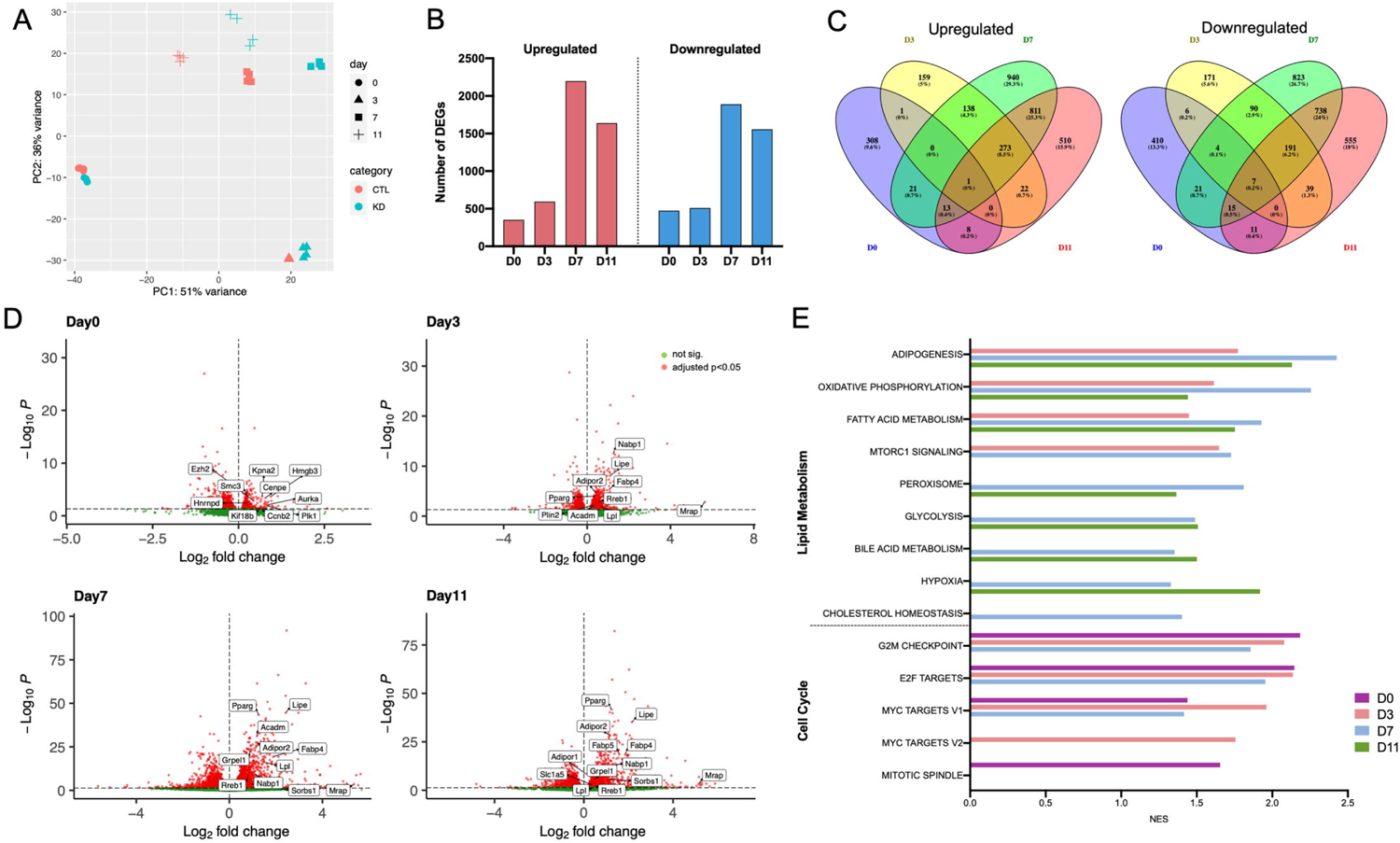
Global transcriptional profiling reveals that *Zbtb9* regulates multiple target genes during adipogenesis. (A) Principle component analysis (PCA) of *Zbtb9* knockdown (KD) or control (CTL) transcriptomes in 3T3-L1 cells at different time points during adipogenesis. (B) Number of differentially expressed genes (DEGs) significantly (FDR < 0.05) up or down regulated upon *Zbtb9* KD in 3T3-L1 cells at D0, D3, D7 and D11. (C) Venn diagrams showing the overlap between DEGs (*Zbtb9*-KD vs. control) upregulated or downregulated at different time points from paired comparisons. (D) Volcano plots showing the DEGs between *Zbtb9*-KD and control 3T3-L1 cells. Highlighted are genes in E2F targets pathway (Day 0) and adipogenesis pathway (Days 3, 7, 11). (E) Gene set enrichment analysis for all DEGs using the Hallmark gene sets. All pathways shown were significantly different in *Zbtb9*-KD vs. control (FDR < 0.05). NES, normalized enrichment score.

**Figure 4.**
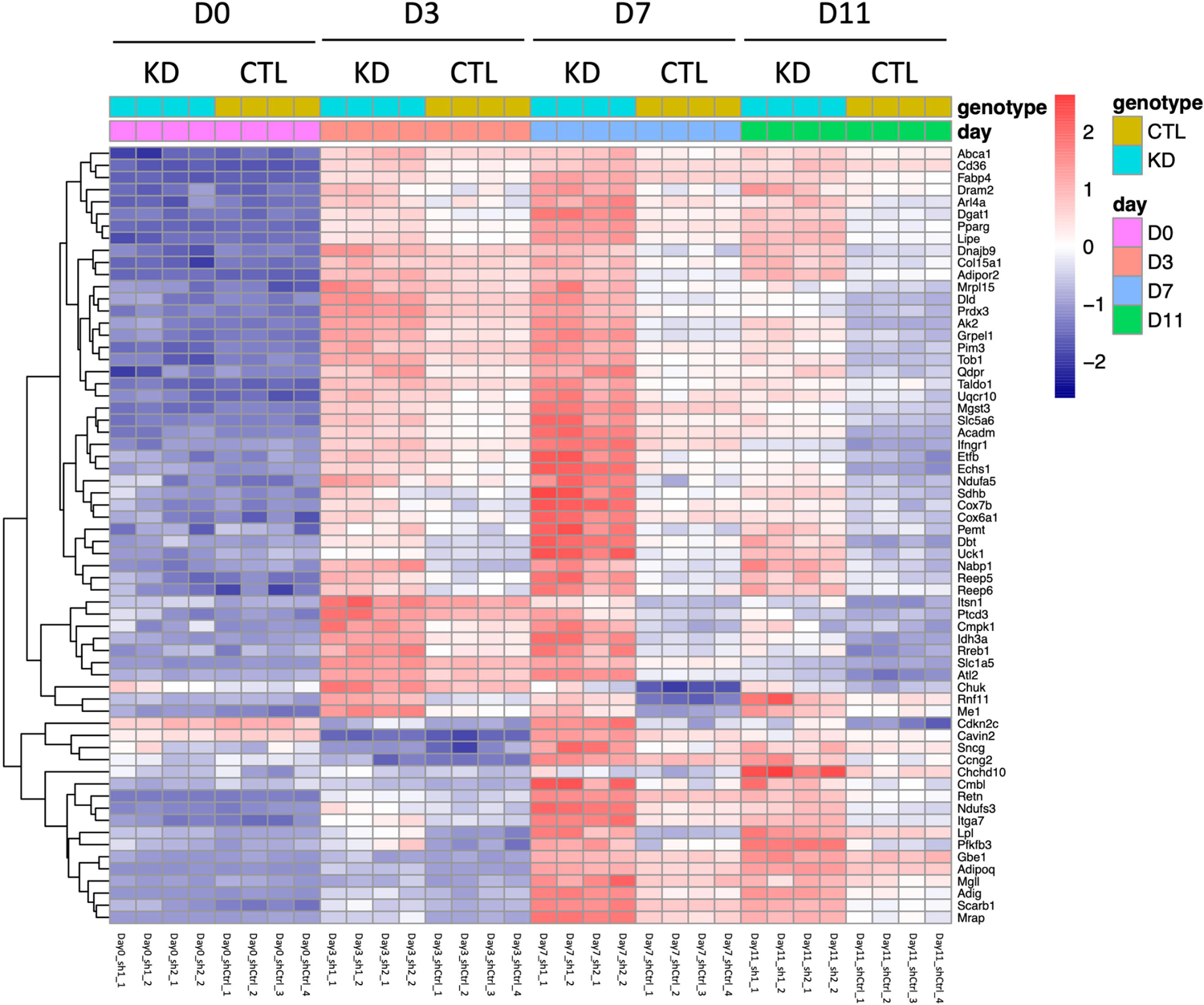
ZBTB9 broadly regulates the expression of genes within the adipogenesis pathway. Heatmap of the adipogenesis pathway in *Zbtb9*-KD vs. control (CTL) 3T3-L1 cells at D0, D3, D7 and D11 based on the RNA-Seq expression profiles. All genes shown in the heatmap were significantly upregulated in KD cells compared to CTL at D3, D7 and D11. sh1_1-2, sh2_1-2, and shCtrl_1-4 indicate individual biological replicates of shRNA#1, shRNA#2, and shCtrl, respectively.

In addition to the downstream effects on adipogenesis likely resulting from increased *Pparg* expression, of interest were the earliest transcriptional differences that led to the upregulation of *Pparg*. For example, at day 0, prior to the induction of adipogenesis, there was no difference in expression levels of *Pparg* between control and *Zbtb9*-KD cells (Fig. 2D) and no upregulation of genes in the adipogenesis pathway (Fig. 3E). Thus, an important question was how *Zbtb9* was driving the increased expression of *Pparg*, one of the earliest and most critical steps in adipogenesis (31). Pathway analysis of the DEGs at day 0, which precedes the increase in adipogenesis gene expression, revealed a significant enrichment of cell cycle-related signaling pathways, including G2M check point, E2F targets, MYC targets and mitotic spindle (Fig. 3E). To identify the transcription factor binding sites in promoter regions of these DEGs in *Zbtb9*-KD cells vs control cells, TransFind (34) was used to predict the transcriptional regulators of the upregulated genes in KD cells relative to control cells at day 0, and identified an enrichment of the canonical E2F binding site in KD cells (Table 1 and Table S1 for detail). The identification of E2F target genes among the most upregulated pathways and enrichment of the E2F consensus binding sites upstream of DEGs was of particular interest given that E2F has previously been shown to directly regulate the expression of *Pparg* both *in vitro* and *in vivo* (35,36).

**Table 1.** Transcription factor motifs enriched in Zbtb9 deficient preadipocytes (Day 0).

| Rank | Transcription Factor | TF matrix | p-value | FDR |
| --- | --- | --- | --- | --- |
| 1 | CNOT3 | V\$CNOT3_01 | 0.000001 | 0.000367 |
| 2 | Staf | V\$STAF_01 | 0.000016 | 0.001818 |
| 3 | Staf | V\$STAF_02 | 0.000016 | 0.001818 |
| 4 | NF-Y | V\$NFY_01 | 0.000016 | 0.001818 |
| **5** | **E2F** | **V\$E2F_Q3** | **0.000016** | **0.001818** |
| **6** | **E2F-1** | **V\$E2F1_Q3** | **0.000084** | **0.008151** |
| 7 | NRF-1 | V\$NRF1_Q6 | 0.000184 | 0.015589 |
| 8 | CREB | V\$CREB_Q2 | 0.00039 | 0.017616 |
| **9** | **E2F** | **V\$E2F_Q4** | **0.00039** | **0.017616** |
| **10** | **E2F** | **V\$E2F_Q6** | **0.00039** | **0.017616** |
| **11** | **E2F** | **V\$E2F_03** | **0.00039** | **0.017616** |
| 12 | ETF | V\$ETF_Q6 | 0.00039 | 0.017616 |
| 13 | AhR | V\$AHR_Q5 | 0.00039 | 0.017616 |
| 14 | MECP2 | V\$MECP2_01 | 0.00039 | 0.017616 |
| **15** | **E2F** | **V\$E2F_02** | **0.000801** | **0.027123** |
| 16 | AP-2 | V\$AP2_Q6 | 0.000801 | 0.027123 |
| 17 | NGFI-C | V\$NGFIC_01 | 0.000801 | 0.027123 |
| 18 | NF-μE1 | V\$NFMUE1_Q6 | 0.000801 | 0.027123 |
| 19 | YY1 | V\$YY1_Q6_02 | 0.000801 | 0.027123 |
| 20 | c-Myb | V\$CMYB_01 | 0.001593 | 0.044925 |
| **21** | **E2F-1** | **V\$E2F1_Q6** | **0.001593** | **0.044925** |
| **22** | **Rb:E2F-1:DP-1** | **V\$E2F1DP1RB_01** | **0.001593** | **0.044925** |
| **23** | **E2F** | **V\$E2F_Q6_01** | **0.001593** | **0.044925** |

\*E2F related motifs are highlighted in bold.

### ZBTB9 modulates RB-E2F signaling to control the early induction of adipogenesis

E2F is a family of transcription factors that play a critical role in early adipogenesis (37,38). Given that *Zbtb9* deficiency causes upregulation of E2F target genes (Fig. 3D, Day 0), we sought to test whether the effects of ZBTB9 on adipogenesis and adipogenic gene expression were mediated by E2F. We first examined the activity of *E2F* in 3T3-L1 preadipocytes utilizing an E2F response element luciferase reporter, and demonstrated a significant increase in *E2F* activity due to *Zbtb9* deficiency (2.2 and 1.8-fold increase with shRNA#1 and #2 respectively), confirming that ZBTB9 does negatively regulate *E2F* activity (Fig. 5A). Next, we tested whether the increase in adipogenic gene expression and adipogenesis due to *Zbtb9* deficiency was dependent on *E2F* activity. Towards this end, *Zbtb9* was knocked down in 3T3-L1 preadipocytes and the cells were induced to undergo adipocyte differentiation, either in the presence or absence of the E2F inhibitor HLM006474 (E2Fi). Consistent with previous results (Fig. 2A-C), *Zbtb9* deficiency again increased adipogenic gene expression and adipogenesis as measured by Oil Red O lipid staining (Fig. 5B-D). However, in the presence of E2Fi, *Zbtb9* deficiency failed to increase levels of *Pparg*, *Adiponectin*, *Glut4*, and *Fabp4*, relative to in the absence of E2Fi when each of these genes was significantly increased (Fig. 5B). Lipid accumulation was also significantly reduced in E2Fi-treated cells compared to DMSO control, as quantified by Oil Red O staining (Fig. 5C-D). These results demonstrate that the effect of ZBTB9 on adipogenesis is dependent on *E2F* activity.

**Figure 5.**
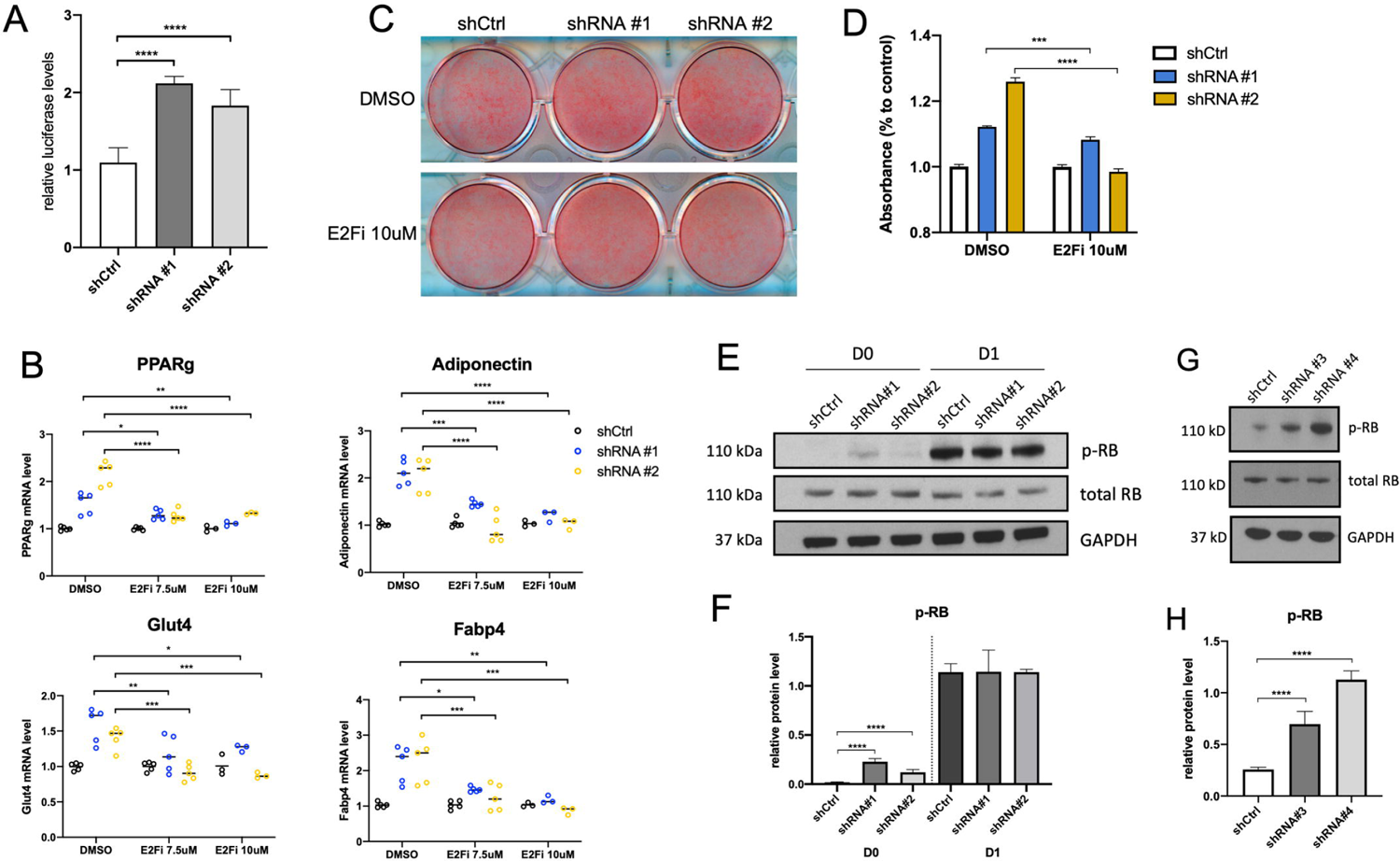
ZBTB9 regulates early induction of adipogenesis through Rb-E2F signaling. (A) An *E2F* consensus reporter plasmid was transfected into *Zbtb9*-KD or control 3T3-L1 preadipocytes. Luciferase activity was measured. (B) *Zbtb9*-KD or control 3T3-L1 preadipocytes were treated with the E2F inhibitor HLM006474 (E2Fi) or DMSO as a control, and differentiated. Adipogenic gene expression was analyzed by qRT-PCR at the end of differentiation (day 11). (C) Oil Red O staining and (D) quantification of these cells treated with E2Fi or DMSO as a control. (E) Western blots of *Zbtb9*-KD and control 3T3-L1 preadipocytes before the initiation of differentiation (D0) or in differentiation medium for one day (D1). Cell lysates were subjected to SDS-PAGE and western blotting for phosphorylated RB (p-RB) and total RB. GAPDH represents a loading control. (F) p-RB protein levels from panel E were quantified by image J. (G) Western blot analysis of p-RB, total RB and GAPDH (loading control) in *ZBTB9*-KD and control human preadipocytes before differentiation. (H) p-RB protein levels from panel G were measured by image J. * p < 0.05, ** p < 0.01, *** p < 0.001, **** p < 0.0001. n=3-5/group.

E2F has been shown to directly regulate the expression of lineage-specifying transcription factors, including PPAR_γ_, both *in vitro* and *in vivo* (35,36), which is independent of its function as a cell cycle regulator. This function of E2F is regulated by the phosphorylation of pocket protein RB (pRB), with pRB dissociating from E2F, enabling the activation and increased transcriptional activity of E2F (39). To test whether ZBTB9 regulates the phosphorylation status of RB, Western blots were performed in control and *Zbtb9* KD cells in both human and mouse preadipocytes. As a control for increased pRB levels, we tested adipogenic induction medium, which is known to induce pRB phosphorylation (38), and which indeed increased phosphorylation of RB as expected (Fig. 5E-F, Day 1). In addition, a clear and reproducible increase in pRB levels was detected in *Zbtb9*-KD mouse preadipocytes compared to control cells, without changing total RB levels (Fig. 5E-F, Day 0, prior to induction of differentiation). In human preadipocytes, *Zbtb9*-KD also increased pRB levels at Day 0 (Fig. 5G-H). The results demonstrated that ZBTB9 regulates adipogenesis via an E2F-dependent mechanism that is associated with increased pRB levels and elevated E2F activity.

## Discussion

Little is known about the cellular function of ZBTB9, although multiple GWAS suggested a role in metabolic disease susceptibility. We now report for the first time that ZBTB9 regulates adipogenesis and adipocyte function, suggesting a possible molecular mechanism underlying the altered risk of obesity and T2D associated with allelic variation near ZBTB9. We demonstrated that ZBTB9 interacts with the PPAR_γ_/RXRα/ZFP407 protein complex in adipocytes and increased PPAR_γ_ activity via the consensus PPAR_γ_/RXRα DNA binding motif. Interestingly, ZBTB9 itself increased the PPAR_γ_ reporter gene expression in HEK293T cells (Fig. 1B), which do not express endogenous PPAR_γ_. Thus, it is possible that ZBTB9 activates PPAR_γ_ target genes by directly binding to the response element with or without interacting with PPAR_γ_. Unlike ZFP407, which regulates PPAR_γ_ activity in the absence of a direct effect on PPAR_γ_ expression, ZBTB9 deficiency reduced PPAR_γ_ gene expression as well as PPAR_γ_ target genes in 3T3-L1 adipocytes (Fig. 1D).

PPAR_γ_ is important for mature adipocyte function (40,41), and crucial for controlling gene networks involved in glucose homeostasis and insulin sensitization (31). PPAR_γ_ transactivation is induced by ligand-dependent and independent mechanisms. Ligand-dependent transactivation is induced by ligand binding to the C-terminal activation function (AF-2) domain (42). PPARs form heterodimers with the RXR and bind to PPAR response elements (PPREs) in enhancers of downstream target genes (43). Binding of a ligand to a PPAR results in the dissociation of a corepressor protein complex and then recruitment of several transcriptional coactivators, some of which are responsible for the modification of histone and chromatin structure to open up DNA for transcription, while others provide linkage to core basal transcriptional machinery (44,45). Many coactivators and corepressors of PPAR_γ_ have been reported over the past two decades, such as *TIF2*, *PGC-1α*, *TRAP220/DRIP205/PBP*, *RIP140*, *NCoR*, *SMRT*, *Sirt1*, and *TAZ* (17,45).

PPAR_γ_ and the coregulators function as multiprotein complexes to activate target gene transcription. Each of the coregulators has its own unique inherent physiological function in lipid and energy metabolism. PPAR_γ_-mediated hormonal or non-hormonal signal transduction regulates cell growth, differentiation, development, metabolism and other important physiological functions. An understanding of the functional significance of individual components of the complicated coregulator complexes in PPAR_γ_ signal transduction pathway will provide multiple drug targets that may fine-tune PPAR_γ_ signaling or better integrate other signaling pathways. Our study shows that ZBTB9 functions as a positive regulator of PPAR_γ_ signaling in mature adipocytes, which adds an important piece to the PPAR_γ_ transcriptional puzzle by discovering a novel protein with its own non-redundant properties in regulating adipogenesis and adipocyte gene expression.

Adipogenesis has been studied extensively *in vivo* and *in vitro* but many questions remain about the exact molecules and mechanisms that govern this process (8,46,47). The growth and expansion of adipocytes and adipose tissue *in vivo* depends on the self-renewal and differentiation of adipose precursor cells (APCs), differentiation into preadipocytes, and finally the differentiation of preadipocytes into mature adipocytes (48). Adipose expansion through adipogenesis can offset the negative metabolic effects of obesity, and the mechanisms and regulators of this adaptive process are now emerging. Adipocyte differentiation involves a temporally regulated set of gene-expression events, and understanding the underlying transcriptional networks is of fundamental importance. PPAR_γ_ is the master regulator of adipogenesis (11,12). Identifying new molecules interacting with PPAR_γ_ will shed light on the function of PPAR_γ_ in adipogenesis. One of the most consequential downstream effects of PPAR_γ_ is the activation of the transcription factor C/EBPα (49). C/EBPα and PPAR_γ_ functionally synergize to fully activate the mature adipocyte program (50,51). Over the past two decades, many factors have been found to regulate adipogenesis. For example, transcription factor ZFP467 suppresses osteogenesis and promotes adipogenesis of the fibroblast-like progenitors by enhancing the expression of C/EBPα (52). Furthermore, KLF5 binds to and activates the *Pparg* promoter, functioning in concert with C/EBPα (53). By contrast, GATA2 and GATA3 inhibit adipogenesis through inhibition of *PPARG* transcription (54).

We further explored the role of ZBTB9 in adipocyte differentiation. Surprisingly, shRNA-mediated *Zbtb9* deficiency in preadipocytes led to an increase in adipogenesis as indicated by both lipid accumulation and adipogenic gene expression. In support of this, pathway analysis of RNA-Seq data revealed significant enrichment of genes in the adipogenesis pathway, with gene expression elevated by *Zbtb9* deficiency shortly after the induction of differentiation (Fig. 4, Day 3). These results were unexpected given the role of ZBTB9 as a positive regulator of PPAR_γ_ in mature adipocytes, as for example is illustrated in Figure 1. In contrast, PPAR_γ_ and its target genes, which are key drivers of adipogenesis, were negatively regulated by ZBTB9 at the early stages of differentiation by an alternative mechanism. Like ZFP467 and KLF5 mentioned above, the identification of ZBTB9 also helps to better define the precise mechanisms in adipogenesis, by recognizing the role of ZBTB9 and demonstrating that it works via the E2F pathway.

Adipogenesis involves two major events: preadipocyte proliferation and adipocyte differentiation (55). *In vitro* studies using 3T3-L1 preadipocyte model have been instrumental in studying this process. Re-entry into cell cycle of growth arrested preadipocytes following differentiation induction is a required initial event occurring during adipogenesis. After several rounds of clonal expansion, cells arrest proliferation again and undergo terminal adipocyte differentiation (56). E2F transcription factors can promote transcriptional activation of genes that encode cell-cycle regulators required for S-phase entry and progression of the cell cycle (57). These events are critical for mitotic clonal expansion, an obligate step in the adipocyte differentiation program (58). E2F also has important metabolic functions beyond the control of the cell cycle (59–61).

For example, E2F1 was demonstrated to be a positive regulator of adipogenesis, by promoting *Pparg* expression or activity, independent of its role as cell cycle regulator (38). Regulation of *Pparg* expression by E2F1 is through direct binding to an E2F-responsive element in the *Pparg* promoter early during adipogenesis (38). Thus, E2Fs represent a link between proliferative signaling pathways, triggering clonal expansion and terminal adipocyte differentiation through regulation of *Pparg* expression.

The pocket protein RB is a major regulator of E2F1 activity. Phosphorylation of RB results in dissociating from E2F, enabling the activation and increased transcriptional activity of E2F (39). RB has an inhibitory role at early stage of adipocyte differentiation, through the formation of a complex including HDAC3 that inhibits PPAR_γ_-dependent gene expression and adipocyte differentiation (62). However, the lack of RB inhibits adipogenesis in 3T3-L1 and MEF cells (63). In addition, mice with a conditional deletion of *Rb* in adipose tissue have increased mitochondrial activity resulting in an increased energy expenditure, which protects them from diet-induced obesity (64). These apparently opposite roles of RB in adipogenesis can be reconciled as during early stage of adipocyte differentiation, cells need to exit cell cycle. In this withdrawal stage RB plays a major role, and positively regulates adipogenesis in a PPAR_γ_-independent manner. Later during differentiation, RB represses PPAR_γ_ activity but the net result is still decreased fat mass in the absence of RB. Thus, the pRB-E2F1 pathway, in which we show that Zbtb9 is a key regulatory molecule, plays an important role in metabolism at different stages of adipogenesis.

The gene expression data in our study shows no difference in expression levels of *Pparg* between *Zbtb9*-KD and control cells at Day 0, which precedes the increase in adipogenic gene expression, but a significant enrichment of E2F target pathway in *Zbtb9*-KD cells compared with control cells at this time point. During this clonal expansion phase of adipocyte differentiation which represents the early stage of adipogenesis, E2F1 regulates the expression of genes implicated in the entry of the cells into cell cycle. E2F1 also promotes *Pparg* expression at this stage, as shown by gene expression results at Day 3 (Fig. 2D, Fig. 4). The luciferase assay indicates increased *E2F* transcriptional activity in *Zbtb9*-KD cells, as compared to control cells, before differentiation. In line with this, RB phosphorylation was also enhanced in *Zbtb9*-KD cells at Day 0, which activated *E2F1*. Our data suggest ZBTB9 regulates adipogenesis via an E2F-dependent mechanism that is associated with increased RB phosphorylation levels and elevated E2F activity.

Based on our data, ZBTB9 plays dual roles in the regulation of PPAR_γ_ signaling in a cell state-dependent manner. Our study provides mechanistic insights into how ZBTB9 regulates early adipogenesis and adipocyte function, identifying a new molecule that may be important in the pathogenesis and treatment of obesity and T2D. Although much remains to be discovered about the underlying molecular mechanism as well as the physiological role of ZBTB9 in adiposity.

### Experimental procedures

#### Cell culture

3T3-L1 cells were passaged and differentiated as previously described (25). Briefly, 3T3-L1 cells were induced to differentiation at day 0 (2 days post confluence) by adding the induction medium, which is the complete culture medium supplemented with the DMI cocktail (1 _μ_M dexamethasone, 0.5 mM 3-isobutyl-1-methylxanthine, and 167 nM insulin, all from Sigma, Saint Louis, MO). At day 3, the induction medium was removed and the maintenance medium (complete culture medium supplemented with 167 nM insulin) was added. At day 7, the maintenance medium was removed, and complete culture medium was added. The cells were harvested for staining or RNA extraction at day 11.

Human preadipocytes were obtained from Sigma (#802S-05A) and plated and cultured with Human Preadipocyte Growth Medium (Sigma #811-500). Once confluent, the cells were subjected to differentiation with Human Preadipocyte Differentiation Medium (Sigma #811D-250) for 12 days. The differentiation medium was refreshed every other day. Cells were harvested for staining or RNA extraction at the end of differentiation.

#### Lentiviral production and infection

Lentiviral particles expressing either a control shRNA (shCtrl: pLKO.1, Sigma-Aldrich, St. Louis, MO, USA) or shRNAs targeting *Zbtb9* (mouse *Zbtb9* shRNA #1: TRCN0000125706; mouse *Zbtb9* shRNA #2: TRCN0000125707; human *ZBTB9* shRNA #3: TRCN0000017185; human *ZBTB9* shRNA #4: TRCN0000017186) were prepared and propagated in HEK293T cells as described previously using the second generation of psPAX2 and pMD2.G packing vectors. After two rounds of lentiviral infection, cells were then cultured and differentiated as previously described (32).

#### Oil Red O staining

To make Oil Red O (ORO) working solution, 20 mL of 0.5% of the ORO stock solution (Sigma #O1391) (in isopropanol) was added to 30 mL of deionized water. Cells were washed with PBS and fixed with 10% formalin at room temperature for 30 min, then stained with the ORO working solution for 15 min. The cells were then rinsed with water three times, and then scanned with an image scanner (EPSON Perfection V600). For quantification, stain was extracted in isopropanol and measured at 492 nm using a BioTek Epoch Reader.

#### RNA Analysis

RNA was collected using QIAshredder and RNeasy mini kit (Qiagen). RNA for qRT-PCR was reversed transcribed using the high capacity cDNA reverse transcription kit without the RNase inhibitor (Applied Biosystems, Carlsbad, CA). Primer sequences are given in Table S2. qRT-PCR was performed in triplicate on a QuantStudio3 Real-time PCR system using the power SYBR Green PCR master mix (Applied Biosystems). The expression level for all genes was calculated using the ΔΔCt method relative to the *Gapdh* control gene. For RNA-Seq analysis, total RNA was isolated as described above with an additional on-column DNase treatment step. RNA quality was determined on the Agilent BioAnalyzer 2100 and all samples had an RNA integrity number score greater than 9.5. Illumina TruSeq sequencing libraries were prepared by Novogene Co., LTD using standard procedures. Samples were run on an Illumina NovaSeq generating an average of 68,511,857 reads per sample. Reads were aligned to the mouse genome (Ensembl m38.102) using TopHat version 2.1.0, SamTools version 1.3.1, and Bowtie2 version 2.2.6 (65–67). Gene expression count tables were generated using HTSeq version 0.6.1 (68) and analyzed for differential expression by DESeq2 version 1.30 (69). A false discovery rate-adjusted *p* value of < 0.05 was considered statistically significant for RNA-Seq analysis. The RNA-Seq expression profiling data are available in the Gene Expression Omnibus series GSE253544.

#### Luciferase reporter assay

To assess *Pparg* activity, HEK293T cells or 3T3-L1 mature adipocytes were transfected with DNA plasmid constructs encoding PPAR_γ_ (Addgene no. 8862), ZBTB9 (MR207329; Origene Technologies, Rockville, MD), or an empty vector control plasmid (pRK5-Myc), together with the *Pparg* target gene luciferase reporter plasmid (PPRE-X3-Tk-luc, Addgene no. 1015), and control plasmid pRL-SV40 encoding Renilla for normalization.

To assess *E2F* activity, *Zbtb9* or control shRNA treated 3T3-L1 preadipocytes were transfected with the *E2F* target gene luciferase reporter plasmid (pGreenFire1_E2F1RE, Addgene no. 112248) and control plasmid pRL-SV40 encoding Renilla for normalization. Luciferase and Renilla were measured 48 hours post-transfection with the Dual-Glo Luciferase Assay System (Promega, Madison, WI).

#### Western blotting

Western blotting was performed and quantitated as described (25). A custom anti-ZFP407 antibody was generated in rabbit against the COOH-terminal 149 amino acids of the mouse ZFP407 protein (Proteintech Group) and has been described previously (25). The anti-ZBTB9 antibody was from Aviva Systems Biology (#ARP31669_P050). The anti-PPAR_γ_ antibody was obtained from Bethyl Laboratories (#A304-461A, Montgomery, TX). The anti-RXRα (#3085), anti-phopho-RB (#8180), anti-RB (#9313) antibodies were from Cell Signaling Technologies (Danvers, MA). Anti-GAPDH was from Proteintech Group (#60004-1, Rosemont, IL). The goat anti-rabbit (31460) and goat anti-mouse (31430) secondary antibodies were from ThermoFisher Scientific (Waltham, MA).

#### Statistical analyses

All data are expressed as the mean ± SEM. Statistical significance was assessed with two-tailed Student’s *t*-Test using GraphPad Prism 8 software. P values < 0.05 were considered statistically significant for all statistical tests.

## Data availability

All RNA-Seq data is available in the Gene Expression Omnibus at accession number GSE253544.

## Supporting information

This article contains supporting information.

## Acknowledgments

This work made use of the High Performance Computing Resource in the Core Facility for Advanced Research Computing at Case Western Reserve University.

## Funding and additional information

This study was supported by the NIDDK grant DK119305 (D.A.B.). The content is solely the responsibility of the authors and does not necessarily represent the official views of the National Institutes of Health.

## Conflict of interest

The authors declare that they have no conflicts of interest with the contents of this article.

---

**Figure S1.** Evolutionary conservation of the mouse and human ZBTB9 protein sequence. Highlighted are the mouse and human ZBTB9 protein BTB domain and 2 zinc finger C2H2 domains.

## The abbreviations used are

PPARγ: peroxisome proliferator-activated receptor-γ

T2D: type 2 diabetes

KD: knockdown

TZD: Thiazolidinedione

RB: retinoblastoma

TAG: triacylglycerol

PGC-1α: PPAR-gamma coactivator-1α

ZFP407: Zinc finger protein 407

RXRα: retinoid X receptor α

BMI: body mass index

PCA: principle component analysis

DEGs: differentially expressed genes

E2Fi: E2F inhibitor

PPREs: PPAR response elements

APCs: adipose precursor cells

ORO: oil red O.

## Notes

### Competing Interest Statement

The authors have declared no competing interest.

