## Supplementary figures and images for "Cell-state dependent regulation of PPAR_γ_ signaling by ZBTB9 in adipocytes"

### Supplemental Figure 1

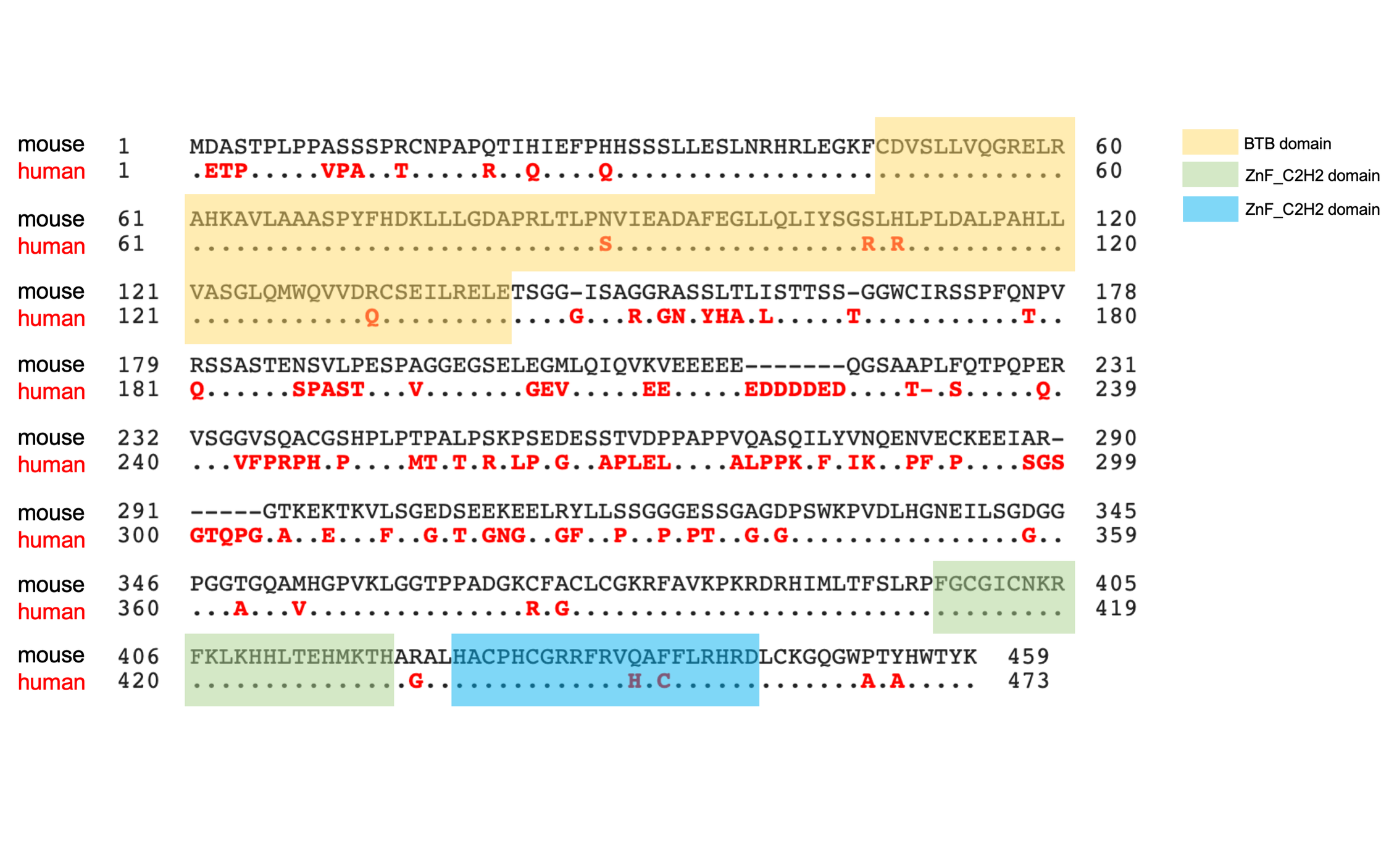
